# Endogenous viral elements are widespread in arthropod genomes and commonly give rise to piRNAs

**DOI:** 10.1101/396382

**Authors:** Anneliek M. ter Horst, Jared C. Nigg, Bryce W. Falk

## Abstract

Arthropod genomes contain sequences derived from integrations of DNA and non-retroviral RNA viruses. These sequences, known as endogenous viral elements (EVEs), have been acquired over the course of evolution and have been proposed to serve as a record of past viral infection. Recent evidence indicates that EVEs can function as templates for the biogenesis of PIWI-interacting RNAs (piRNAs) in some mosquito species and cell lines, raising the possibility that EVEs may function as a source of immunological memory in these organisms. However, whether EVEs are capable of acting as templates for piRNA production in other arthropod species is unknown. Here we used publically available genome assemblies and small RNA sequencing datasets to characterize the repertoire and function of EVEs across 48 arthropod genomes. We found that EVEs are widespread in arthropod genomes and primarily correspond to unclassified ssRNA viruses and viruses belonging to the Rhabdoviridae and Parvoviridae families. Additionally, EVEs were enriched in piRNA clusters in a majority of species and we found that production of primary piRNAs from EVEs is common, particularly for EVEs located within piRNA clusters. While we found evidence suggesting that piRNAs mapping to a number of EVEs are produced via the ping-pong cycle, potentially pointing towards a role for EVE-derived piRNAs during viral infection, limited nucleotide identity between currently described viruses and EVEs identified here likely limits the extent to which this process plays a role during infection with known viruses.

## BACKGROUND

Arthropods play key roles in terrestrial and aquatic ecosystems by pollinating plants, aiding in plant seed dispersal, controlling populations of other organisms, functioning as food sources for other organisms, and cycling nutrients [1, 2]. Besides their important contributions to maintaining ecosystem stability, some arthropods are also known to serve as vectors for human, animal, and plant pathogens [3, 4]. During arthropod-mediated transmission of many plant- and animal-infecting viruses, the virus replicates inside the arthropod vector, thus the vector serves as one of at least two possible hosts for these viruses [3, 4]. Additionally, arthropods are subject to infection by arthropod specific viruses that are not transmitted to new hosts [5]. Elucidating the antiviral mechanisms arthropods use to combat viral infection is an important area of research, as a greater understanding of arthropod immunity may lead to new strategies for the control of arthropod-transmitted viruses.

RNA interference (RNAi) is the primary antiviral mechanism in arthropods and relies on three classes of small RNAs (sRNAs) [6, 7]. The small interfering RNA (siRNA) pathway is the most important branch of RNAi for combating viral infection in arthropods and this pathway relies on the production of primarily 21 nt siRNAs via cleavage of viral double-stranded RNA [7]. siRNAs associate with argonaute proteins to direct a multi-protein effector complex known as the RNA-induced silencing complex to the viral RNA, resulting in endonucleolytic cleavage of target RNA [7]. The micro RNA (miRNA) pathway relies primarily on inhibition of translation via imperfect base pairing between miRNAs and viral RNAs, but miRNAs can also direct cleavage of target RNA if there is sufficient complementarity between the miRNA and the target RNA [7]. A third branch of RNAi directed by PIWI-interacting RNAs (piRNAs) was discovered more recently and has been implicated as a component of antiviral defense in mosquitoes, but not in *Drosophila melanogaster* [8, 9].

The primary role of the piRNA pathway is control of transposable elements in animal germ cells and studies in *D. melanogaster* have revealed two models for piRNA biogenesis: the primary pathway and the ping-pong cycle (secondary pathway) [10]. In the primary pathway, 24-32 nt primary piRNAs with a strong bias for uracil as the 5’-most nucleotide (1U bias) are produced from endogenous transcripts derived from regions of the genome denoted as piRNA clusters. piRNA clusters contain a high load of sequences derived from transposable elements and generally primary piRNAs are antisense to RNAs produced by corresponding transposable elements [11]. During the ping-pong cycle in *D. melanogaster*, antisense primary piRNAs guide the PIWI family argonaute protein Piwi to transposable element RNA, resulting in endonucleolytic cleavage of transposable element RNA exactly 10 nt downstream from the 5’ end of the guiding primary piRNA [10]. Cleaved transposable element RNA is subsequently processed into sense secondary piRNAs with a bias for adenine as the 10th nucleotide from the 5’ end (10A bias). Secondary piRNAs are then loaded onto Aubergine, another PIWI family argonaute protein, and direct cleavage of endogenous transcripts derived from piRNA clusters, resulting in the production of additional primary piRNAs [10]. Thus, the ping-pong cycle serves to amplify the post-transcriptional silencing activity of the piRNA pathway in response to active transposable elements. Interestingly, the PIWI family has undergone expansion in mosquitoes and it is now clear that the mechanisms responsible for generating virus-derived piRNAs in these organisms are distinct from the canonical piRNA pathway used to combat transposable element activity [12, 13]. Key to the novel piRNA pathway seen in mosquitoes is the biogenesis of primary piRNAs directly from exogenous viral RNA without the need for primary piRNAs derived from endogenous sequences [13].

Recent studies have revealed that the genomes of some eukaryotic species contain sequences derived from integrations of DNA and non-retroviral RNA viruses [14-18]. These sequences are known as endogenous viral elements (EVEs) and are proposed to serve as a partial record of past viral infections [15]. Moreover, a number of studies have demonstrated that EVEs are present within piRNA clusters and serve as sources of piRNAs in certain mosquito species and cell lines, raising the possibility that EVEs may participate in an antiviral response against exogenous viruses via the canonical piRNA pathway [14, 15, 18]. While EVEs have been reported in a number of other arthropod species, their potential involvement with the piRNA pathway remains unclear. Here we sought to expand knowledge of EVEs and their role in the piRNA pathway beyond mosquito species. To this end we performed a comprehensive analysis to characterize the abundance, diversity, distribution, and function of EVEs across all arthropod species with sequenced genomes for which there are corresponding publically available sRNA sequencing data. Our results reveal that, as has been observed in mosquitoes, EVEs are abundant in arthropod genomes and many EVEs produce primary piRNAs. Additionally, we found evidence suggesting that piRNAs mapping to a number of EVEs are produced via the ping-pong cycle, potentially pointing towards a role for EVE-derived piRNAs during viral infection. However, limited nucleotide identity between currently described viruses and EVEs identified here likely limits the extent to which this process plays a role during infection with known viruses.

## MATERIALS AND METHODS

### Data Collection

A list of currently sequenced arthropod genomes was retrieved from the 5,000 insect genome project [19]. Genome sequences were then retrieved from GenBank for all species with sRNA sequencing data available in the NCBI sequence read archive (SRA). The accession numbers for all genome assemblies analyzed are available in additional file 1. For each arthropod species, a representative collection of available sRNA datasets was retrieved from the NCBI SRA and the datasets were combined for analysis. The accession numbers of sRNA datasets used for each species are available in additional file 2.

### Identification of EVEs

To identify EVEs in arthropod genomes, we created a BLAST database containing all ssRNA, dsRNA, and ssDNA virus protein sequences available in GenBank. We did not include dsDNA viruses in our analysis due to the difficulty in unambiguously characterizing dsDNA-viral sequences to be of viral origin due to the frequency of horizontal gene transfer between dsDNA viruses and their hosts, and between dsDNA viruses and transposable elements. For each arthropod species, we searched for matches to our viral protein database genome wide using BLASTx with an evalue of 0.001. As reported previously, we found that a large number of putative EVEs identified by this process could not be unambiguously classified as viral sequences due to homology with eukaryotic, bacterial, or archaeal sequences [15]. Such artifacts were initially filtered out of the dataset using custom scripts to extract the genomic nucleotide sequence corresponding to each BLASTx hit (i.e. putative EVEs) and then performing a reverse BLASTx search with these nucleotide sequences against the *D. melanogaster* proteome (Uniprot proteome ID UP000000803) with an evalue of 0.001. Any putative EVEs with a BLASTx hit against the *D. melanogaster* proteome were subsequently removed from analysis. Following this initial filter, the viral proteins corresponding to each putative EVE were compared to the non-redundant protein database by BLASTp and the results were screened manually. If the putative EVE corresponded to a portion of the viral protein possessing a non-viral BLAST hit or a conserved domain with a non-viral lineage (ex. zinc finger domains) then it was removed from the dataset.

Custom python scripts were used to remove duplicate and overlapping EVEs. When two EVEs overlapped, the EVE with the higher BLASTx score was retained. An EVE was defined as one continuous BLASTx hit. Custom python scripts were then used to assign a viral family to each EVE.

### Identification of EVEs in piRNA clusters

Adapter sequences were removed from the sRNA datasets with Cutadapt (version 1.16) using the default settings with the exception that reads as short as 18 nt were retained [20]. After trimming, all the sRNA datasets for each species were concatenated into one dataset per species. These concatenated sRNA datasets were used for all further analysis. piRNA clusters were defined with proTRAC (v2.3.1) using the default settings with the following exceptions: sliding window size = 1000, sliding window increment = 500, threshold clustersize = 1500, and threshold-density p-value = 0.1 [21]. We identified EVEs within piRNA cluster sequences obtained with proTRAC as described above for identification of EVEs genome wide. Custom python scripts were then used to remove any EVEs from the genome wide EVE list that were present in the piRNA cluster EVE list. If an EVE was partially inside and partially outside a piRNA cluster, it was marked as residing outside the piRNA cluster.

### Small RNA mapping and piRNA identification

Concatenated sRNA reads were mapped to arthropod genomes with bowtie (version 1.1.2), using the default settings [22]. Individual BAM files corresponding to each EVE were then generated using samtools based on the genomic coordinates of each EVE and sRNAs mapping to each EVE were extracted from these BAM files using bedtools [23, 24]. Custom python scripts were used to calculate whether an EVE served as a source of primary piRNAs. This was defined as a significant 1U bias (p < .001, cumulative binomial distribution) for 24-32 nt sRNAs mapping to one strand of the EVE. Unlike some other previously described approaches, our analysis examined 1U biases on either strand individually and did not require primary piRNAs to be derived from the antisense strand with respect to the coding potential of the EVEs.

To determine whether sRNAs mapping to each EVE possessed a significant ping-pong signature we first used custom python scripts to calculate whether 24-32 nt sRNAs mapping to each EVE possessed a significant 1U bias as described above. If a 1U bias was observed for sRNAs mapping to one strand, we determined whether 24-32 nt sRNAs mapping to the opposite strand possessed a significant 10A bias (p < .001, cumulative binomial distribution). We then used signature.py to calculate a ping-pong Z-score for 24-32 nt sRNAs mapping to each EVE [25]. sRNAs mapping to each EVE were classified as possessing a significant ping-pong signature if we observed significant 1U and 10A biases for 24-32 nt sRNAs mapping to opposing strands and if the ping-pong Z-score was ≥ 3.2905 (which corresponds to p-value of 0.001 for a two-tailed hypothesis).

### Calculation of nucleotide identities

For each EVE sharing ≥ 75% deduced amino acid identity with its closest viral hit by BLASTx, we retrieved the nucleotide sequence of the EVE using the genomic coordinates. Each EVE nucleotide sequence was then compared to the NCBI non-redundant nucleotide collection via BLASTn. The nucleotide identity obtained via BLASTn was reported. The viral sequences identified by BLASTn were not always the same sequences initially identified by BLASTx (ex. BLASTn identified a strain represented in the non-redundant nucleotide collection, but not represented in the non-redundant protein database). Thus, for calculation of deduced amino acid identities between EVEs and the viral sequences identified by BLASTn, the nucleotide sequences present in the BLASTn alignments were translated and compared via BLASTx

## RESULTS

### EVEs are commonly found within arthropod genomes

We began by identifying all arthropod species for which there are both publically available genome assemblies and sRNA sequencing datasets. We then created a custom database comprised of all ssDNA, ssRNA, and dsRNA viral protein sequences available in GenBank and used this database to identify putative EVEs genome wide in each arthropod genome via BLASTx. As reported previously, we found that a large number of putative EVEs could not be unambiguously classified as viral due to homology with eukaryotic, bacterial, or archaeal sequences [15]. We removed the majority of putative EVEs homologous to eukaryotic sequences via reverse BLAST searches against the *D. melanogaster* proteome. The remaining putative EVEs were then filtered manually. Ultimately, we identified 4,061 EVEs within the genomes of 48 arthropod species (Table 1 & Additional files 3-4). With the exception of *Sarcoptes scabiei*, we found at least one EVE in each arthropod genome.

**Table 1.**
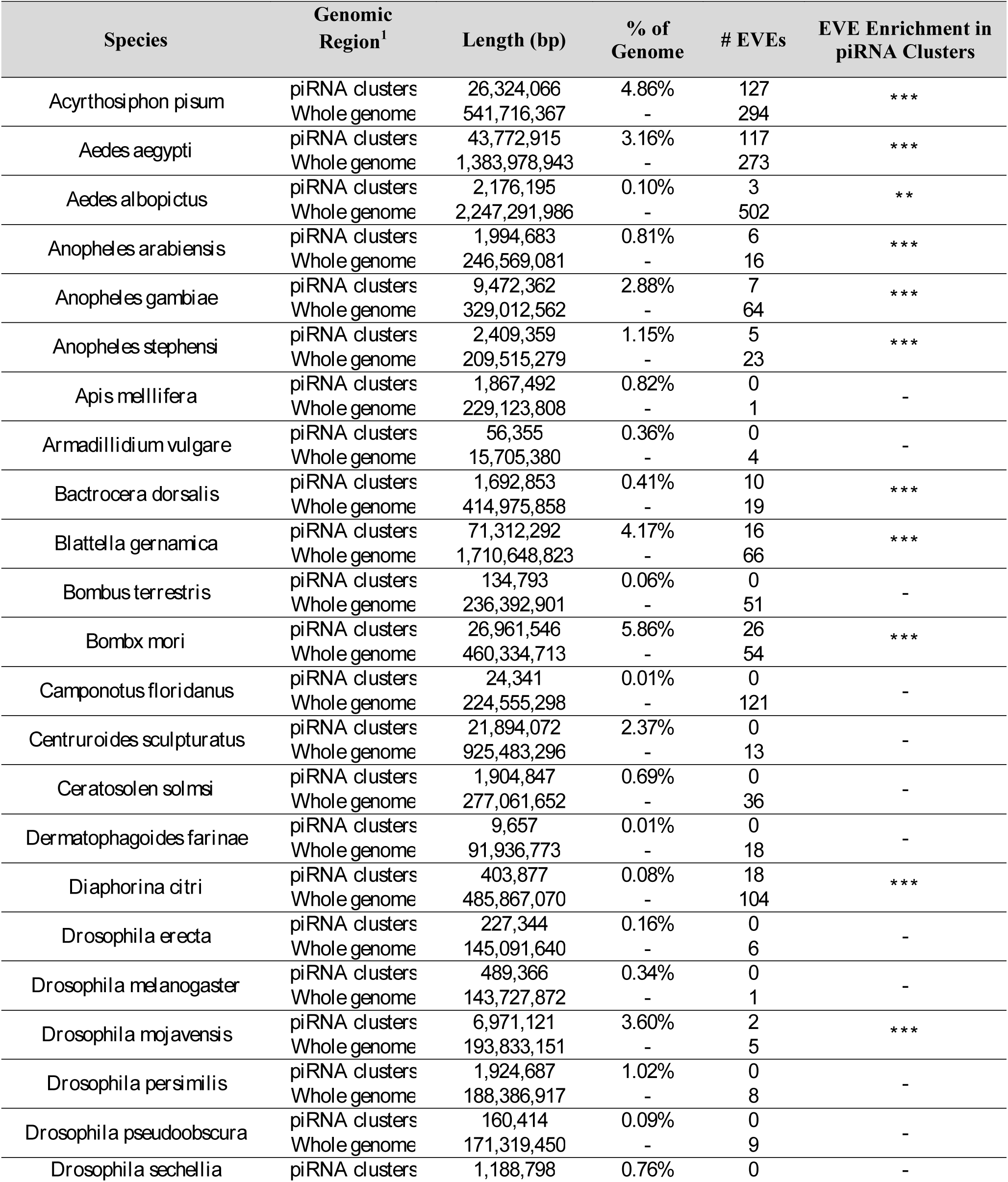
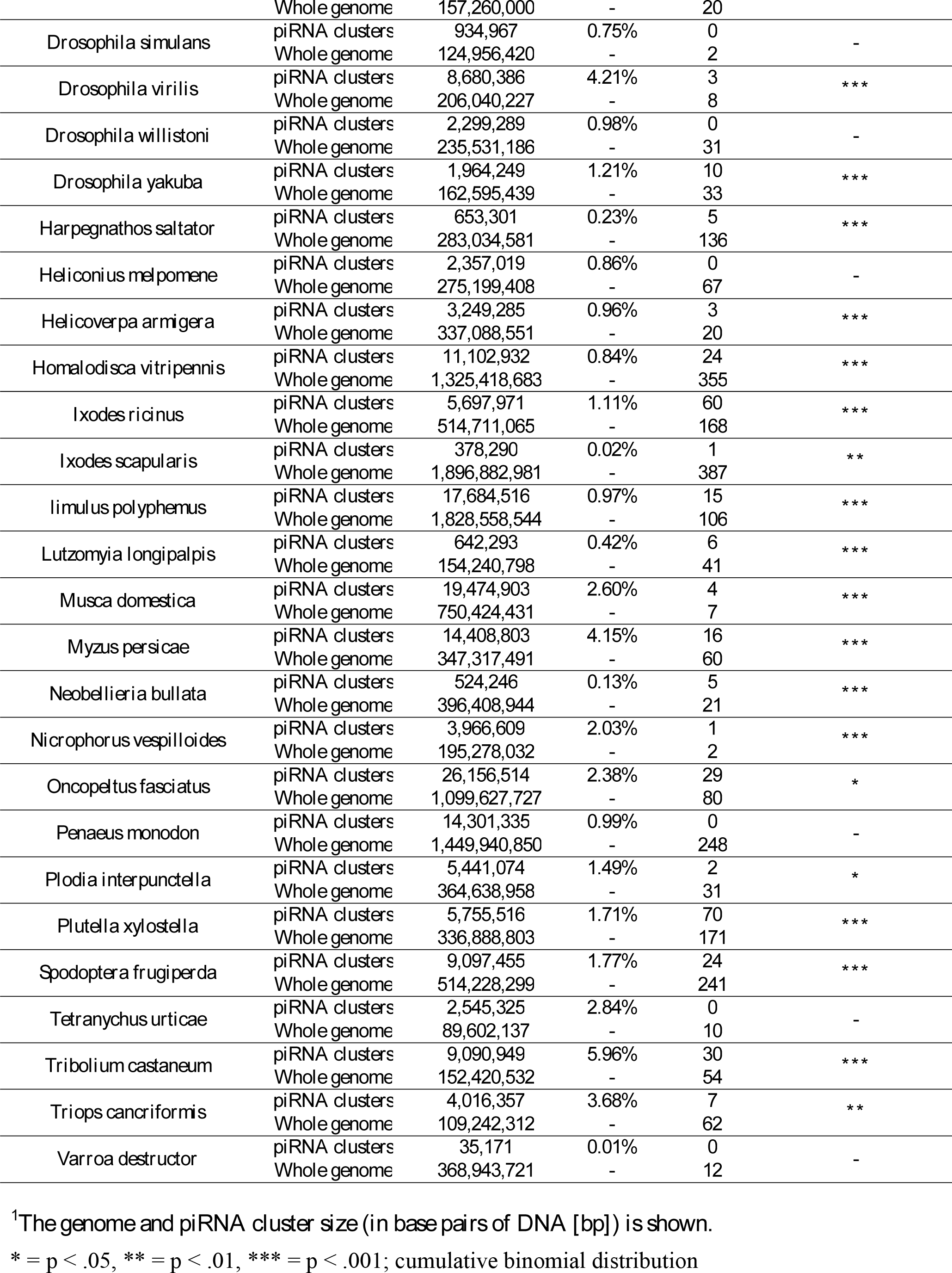
Enrichment of EVEs in piRNA clusters

The 48 arthropod genomes analyzed here contained a median of 1.28 EVEs/10^7^ bp. Notable exceptions include *Apis mellifera* and *Musca domestica,* the genomes of which contained 4.36 EVEs/10^9^ bp and 9.33 EVEs/10^9^ bp, respectively. Interestingly, the ten *Drosophila* sp. genomes analyzed also contained a relatively low number of EVEs with a median of 4.19 EVEs/10^8^ bp. With 5.68 EVEs/10^7^ bp, the *Triops cancriformis* genome contained the largest number of EVEs relative to the size of the genome (Table 1).

### EVEs are enriched in piRNA clusters in a majority of species

Previous studies have pointed towards a potential role for EVE-derived piRNAs in antiviral responses, and EVEs are enriched in piRNA clusters in *Aedes albopictus* and *Aedes aegypti* [14, 25]. Thus, we used publically available sRNA datasets to define piRNA clusters in the arthropod genomes using proTRAC [21]. To increase the coverage and diversity of sRNAs used for this analysis, we combined representative collections of the available sRNA datasets for each species (Additional file 2). We then classified the EVEs into EVEs within piRNA clusters and EVEs outside piRNA clusters (Table 1 & Additional files 3-4). We found that 30 out of 48 arthropod genomes contained EVEs within piRNA clusters and that EVEs were enriched in piRNA clusters in 28 of these species (Table 1). The median deduced amino acid identity shared between EVEs and their closest BLASTx hit was 34.0% for EVEs in piRNA clusters and 34.3% for EVEs outside piRNA clusters. We found that deduced amino acid identity was significantly higher for piRNA cluster resident EVEs in four species and significantly lower in three species (Fig. 1a). Interestingly, we found that when all species are considered, EVEs in piRNA clusters are significantly longer than EVEs outside piRNA clusters (p = .000101, two-tailed T-test). On an individual species level, EVEs were significantly longer within piRNA clusters in seven species and significantly lower in one species (Fig. 1b).

**Fig. 1.**
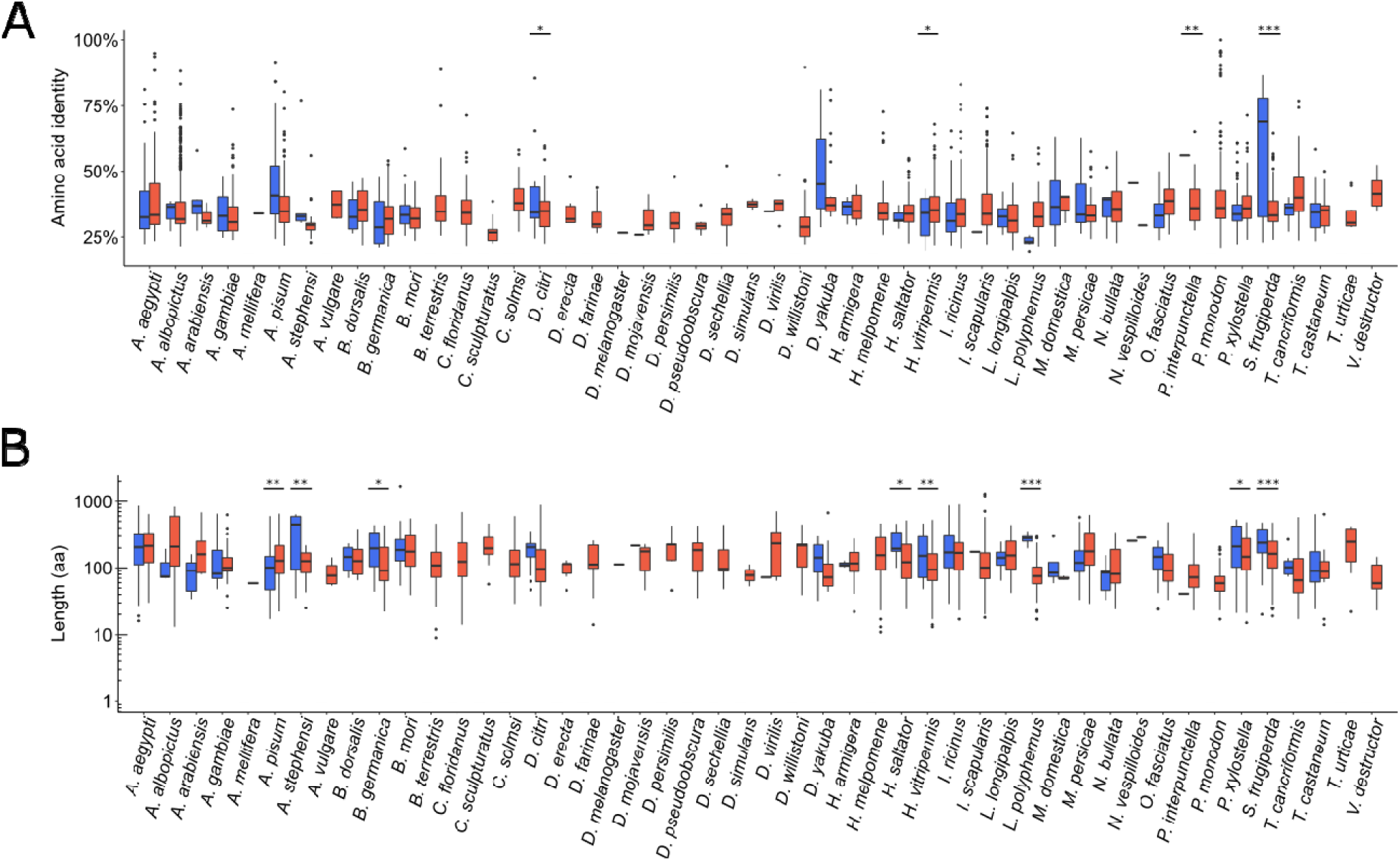
(A) Distribution of amino acid identities between translated EVEs and their closest viral BLASTx hit for the respective arthropod species listed. (B) Distribution of translated EVE lengths in amino acids for the respective arthropod species listed. Blue = EVEs in piRNA clusters, red = EVEs outside piRNA clusters. * = p < .05, ** = p < .01, *** = p < .001; unpaired T-test

### EVEs corresponding to unclassified viruses and viruses belonging to the *Rhabdoviridae* and *Parvoviridae* families predominate both within and outside piRNA clusters

Genome wide, we identified EVEs corresponding to viruses belonging to 54 different viral families (Additional files 5-6). Both within and outside piRNA clusters, unclassified viruses and viruses belonging to the *Rhabdoviridae* and *Parvoviridae* families comprised over 70% of all EVEs (Fig. 2). Interestingly a plurality of EVEs corresponded to viruses possessing negative sense ssRNA genomes (data not shown).

**Fig. 2.**
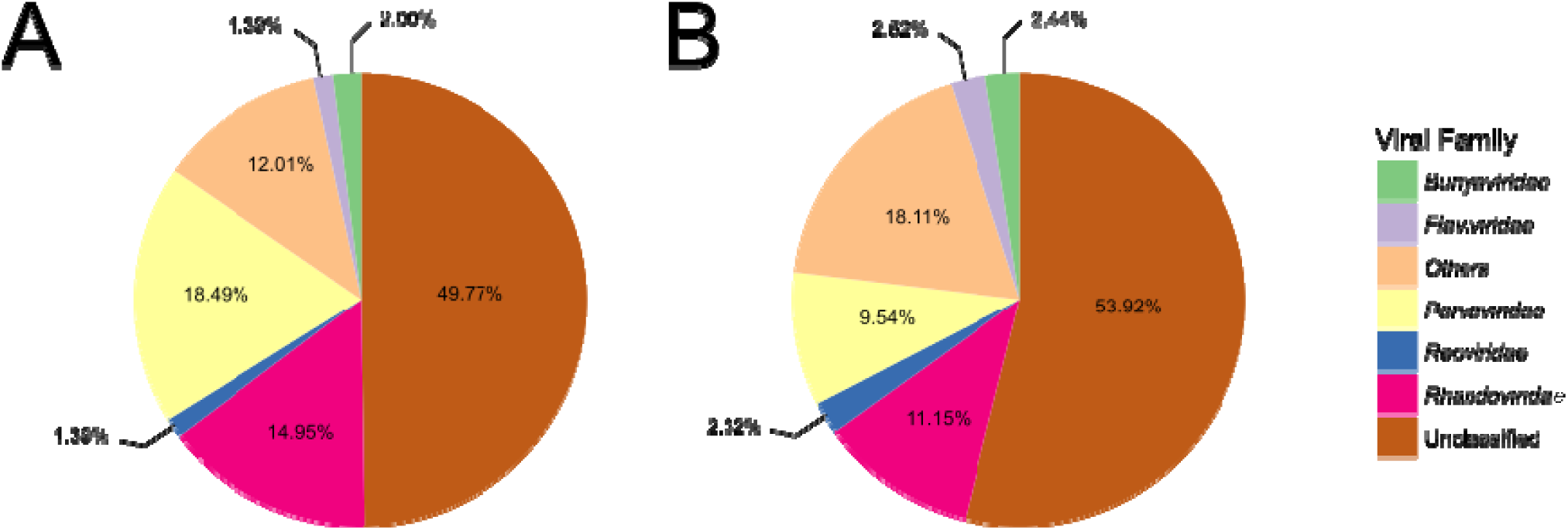
The most common viral families corresponding to EVEs found in arthropod genomes within piRNA clusters (A) or outside piRNA clusters (B). Complete lists of viral families corresponding to EVEs found within arthropod genomes are available in the supplementary information (Additional files 5-6).

Whitfield et al. reported the presence of EVEs corresponding to viruses belonging to the *Closteroviridae* and *Bromoviridae* families within the genome of *A. aegypti*-derived Aag2 cells [15]. This is somewhat unexpected, as these families are comprised solely of viruses that do not infect *A. aegypti*, but only infect plants. These viruses are transmitted by their respective insect vectors in a non-circulative manner [3]. In agreement with these findings, we also identified a number of EVEs corresponding to viruses of the *Closteroviridae* and *Bromoviridae* families, as well as several other families comprised of viruses not known to replicate outside their plant hosts including *Geminiviridae, Nanoviridae, Luteoviridae, Potyviridae, Secoviridae, Tombusviridae*, and *Virgaviridae* (Additional files 5-6).

### Primary piRNA production from EVEs is widespread, but nucleotide identity between EVEs and known viruses is low

Previous studies have revealed that EVEs serve as a templates for piRNA production in *A. aegypti, A. albopictus*, and *Culex quinquefasciatus* [14, 15, 26], however, it is unclear whether piRNAs are produced from EVEs in non-mosquito arthropod species. We examined the sRNAs mapping to each EVE for the characteristics of primary piRNAs (i.e. lU bias for sRNAs 24-32 nt in length). Some previous studies have assessed primary piRNA production from EVEs by measuring 1U biases only for sRNAs mapping antisense with respect to the coding region of the EVE (based on comparison to the corresponding virus). However, primary piRNAs could theoretically be produced from precursor transcripts derived from either genomic strand. Thus, we evaluated 1U biases for 24-32 nt sRNAs mapping either sense or antisense to each EVE. Biases were calculated using a cumulative binomial distribution and deemed significant when p *<* 0.001. We found that the vast majority (81.4%) of EVEs within piRNA clusters served as sources of primary piRNAs. Outside of piRNA clusters, only 35.7% of EVEs served as sources of primary piRNAs. These results indicate that, as in *A. albopictus, A. aegypti,* and *C. quinquefasciatus*, primary piRNAs are frequently derived from EVEs. piRNA production from EVEs was particularly common in *A. aegypti, A. albopictus, Acyrthosiphon pisum, Anopheles stephensi, Bactrocera dorsalis*, and *Nicrophorus vespilloides,* with over 75% of EVEs genome wide serving as templates for primary piRNA biogenesis in these species (Fig. 3). piRNAs were not detected from EVEs in 14 species. Of these, 11 species did not possess EVEs within piRNA clusters.

**Fig. 3.**
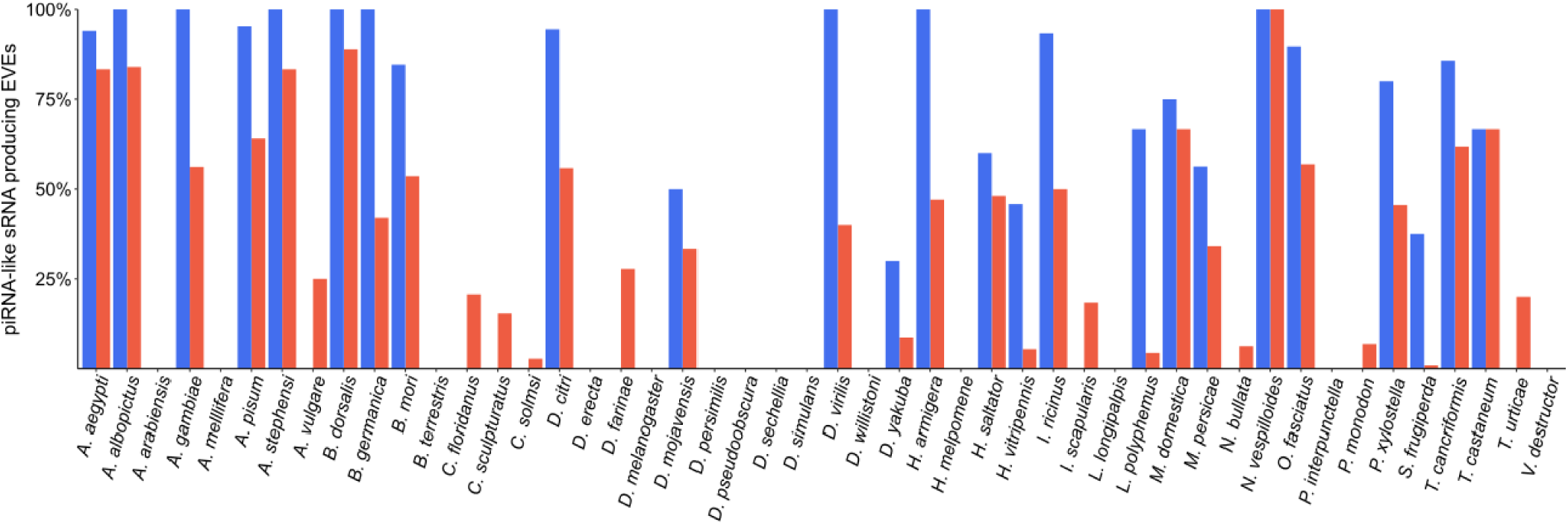
Percent of EVEs producing primary piRNAs for each arthropod species. Blue = EVEs in piRNA clusters, red = EVEs outside piRNA clusters. Primary piRNA production from an EVE was defined as a significant (p < .001, cumulative binomial distribution) 1U bias for 24-32 sRNAs mapping to the EVE.

Given what is known regarding the effects of sequence identity on piRNA-directed cleavage, targeting of exogenous viruses by EVE-derived piRNAs likely requires extensive complementarity between EVEs and corresponding viruses [27, 28]. To elucidate the targeting potential of EVEs identified here, we extracted nucleotide sequences for all EVEs with ≥ 75% deduced amino acid identity with their closest viral hit via BLASTx. We then used BLASTn to calculate the nucleotide identities between these EVEs and corresponding viruses. We found only 13 EVE-virus pairs with nucleotide identity ≥ 90%, and only 17 pairs with at least one ≥ 20 nt region of perfect identity (Table 2). These results indicate that, with the exception of a small number of EVE-virus pairs, nucleotide identity between EVEs and known viruses is likely too low to permit targeting of known viruses by EVE-derived piRNAs in the species analyzed here.

**Table 2.** Nucleotide identity between select EVEs and the closest known virus

| Arthropod Species | Viral Species | Length<br>(nt) | Identity<br>(aa) | Identity<br>(nt) | Longest<br>Identical Regio |
| --- | --- | --- | --- | --- | --- |
| *Penaeus monodon* | Penaeus stylirostris penstyldensovirus 2 | 73 | 100 | 100 | 73 |
| *Penaeus monodon* | IHHNV | 67 | 100 | 100 | 67 |
| *Penaeus monodon* | IHHNV | 48 | 100 | 100 | 48 |
| *Penaeus monodon* | IHHNV | 95 | 100 | 100 | 95 |
| *Penaeus monodon* | IHHNV | 239 | 100 | 100 | 239 |
| *Penaeus monodon* | Decapod penstyldensovirus 1 | 68 | 100 | 100 | 68 |
| *Penaeus monodon* | Penaeus stylirostris penstyldensovirus 2 | 212 | 100 | 99 | 187 |
| *Penaeus monodon* | IHHNV | 380 | 99 | 99 | 177 |
| *Penaeus monodon* | Penaeus stylirostris densovirus | 71 | 100 | 99 | 38 |
| *Penaeus monodon* | IHHNV | 284 | 99 | 99 | 268 |
| *Aedes aegypti* | Cell fusing agent virus | 239 | 96 | 98 | 139 |
| *Penaeus monodon* | IHHNV | 250 | 99 | 98 | 114 |
| *Aedes aegypti* | Cell fusing agent virus | 188 | 94 | 96 | 72 |
| *Triops cancriformis* | SACDV-21 | 63 | 76 | 89 | 22 |
| *Diaphorina citri* | Diaphorina citri densovirus | 617 | 86 | 87 | 46 |
| *Acyrtosiphon pisum* | Dysaphis plantaginea densovirus | 71 | 91 | 87 | 18 |
| *Acyrtosiphon pisum* | Dysaphis plantaginea densovirus | 55 | 82 | 85 | 14 |
| *Acyrtosiphon pisum* | Dysaphis plantaginea densovirus | 55 | 82 | 85 | 14 |
| *Acyrtosiphon pisum* | Dysaphis plantaginea densovirus | 104 | 76 | 85 | 21 |
| *Acyrtosiphon pisum* | Myzus persicae nicotianae densovirus | 165 | 83 | 84 | 17 |
| *Aedes aegypti* | Liao ning virus | 467 | 90 | 82 | 32 |
| *Aedes aegypti* | Grenada mosquito rhabdovirus 1 | 56 | 94 | 82 | 11 |
| *Drosophila yakuba* | Drosophila melanogaster sigma virus | 110 | 83 | 81 | 16 |
| *Aedes aegypti* | North Creek virus | 47 | 80 | 79 | 14 |
| *Drosophila yakuba* | Drosophila melanogaster sigma virus | 92 | 83 | 79 | 13 |
| *Ixodes ricinus* | Jingmen tick virus | 1125 | 84 | 77 | 20 |
| *Spodoptera frugiperda* | Spodoptera frugiperda rhabdovirus | 57 | 84 | 77 | 8 |
| *Drosophila yakuba* | Drosophila melanogaster sigma virus | 203 | 81 | 75 | 14 |
| *Aedes albopictus* | Aedes flavivirus | 1716 | 89 | 74 | 17 |
| *Aedes albopictus* | Kamiti River virus | 420 | 81 | 74 | 16 |
| *Anopheles stephensi* | Hubei virga-like virus 21 | 87 | 90 | 74 | 18 |
| *Drosophila willistoni* | Hubei diptera virus 17 | 115 | 89 | 74 | 16 |
| *Aedes albopictus* | Australian Anopheles totivirus | 323 | 83 | 73 | 14 |
| *Spodoptera frugiperda* | Spodoptera frugiperda rhabdovirus | 541 | 89 | 72 | 11 |
| *Aedes aegypti* | Australian Anopheles totivirus | 372 | 82 | 71 | 13 |
| *Spodoptera frugiperda* | Spodoptera frugiperda rhabdovirus | 695 | 84 | 71 | 17 |
| *Aedes aegypti* | Australian Anopheles totivirus | 616 | 81 | 70 | 14 |
| *Aedes aegypti* | Tongilchon virus 1 | 131 | 84 | 70 | 11 |
| *Ixodes ricinus* | Deer tick mononegavirales-like virus | 2454 | 77 | 70 | 17 |
| *Spodoptera frugiperda* | Spodoptera frugiperda rhabdovirus* | 1137 | 79 | 69 | 17 |
| *Spodoptera frugiperda* | Spodoptera frugiperda rhabdovirus | 841 | 79 | 69 | 11 |

IHHNV = Infectious hypodermal and hematopoietic necrosis virus, SACDV = Sewage- associated circular DNA virus.
\* The *S. frugiperda* genome assembly contains seven duplications of this EVE

### sRNAs mapping to some EVEs show evidence of production via the ping-pong cycle

While we found that nucleotide identity between EVEs and known viruses is generally low, which likely precludes induction of the ping-pong cycle by EVE-derived piRNAs upon infection with known viruses, currently described virus species are thought to represent only a small fraction of total viral diversity, particularly for arthropod-infecting viruses [29]. Thus, there is a possibility that EVE-derived piRNAs could target undescribed viruses and the presence of ping-pong signatures in piRNAs mapping to EVEs would be one indication of the possible functionality of EVE-derived piRNAs. After defining EVEs that produced primary piRNAs (Fig. 3), we assessed whether 24-32 nt sRNAs mapping to these EVEs possessed significant ping-pong signatures. We defined a significant ping-pong signature as 1U and 10A biases for 24-32 nt sRNAs mapping to opposing strands and a ping-pong Z-score of ≥ 3.2905. We found that sRNAs mapping to 3.4% of all EVEs displayed evidence of production via the ping-pong cycle with 20 species possessing at least one EVE displaying evidence of ping-pong dependant piRNA production (Table 3). This number was slightly higher for EVEs within piRNA clusters (5.37%) than for EVEs outside piRNA clusters (3.05%). While further experiments are necessary, we propose that one explanation for the observed ping-pong signatures could be infection with undescribed viruses corresponding to primary piRNA-producing EVEs.

**Table 3.**
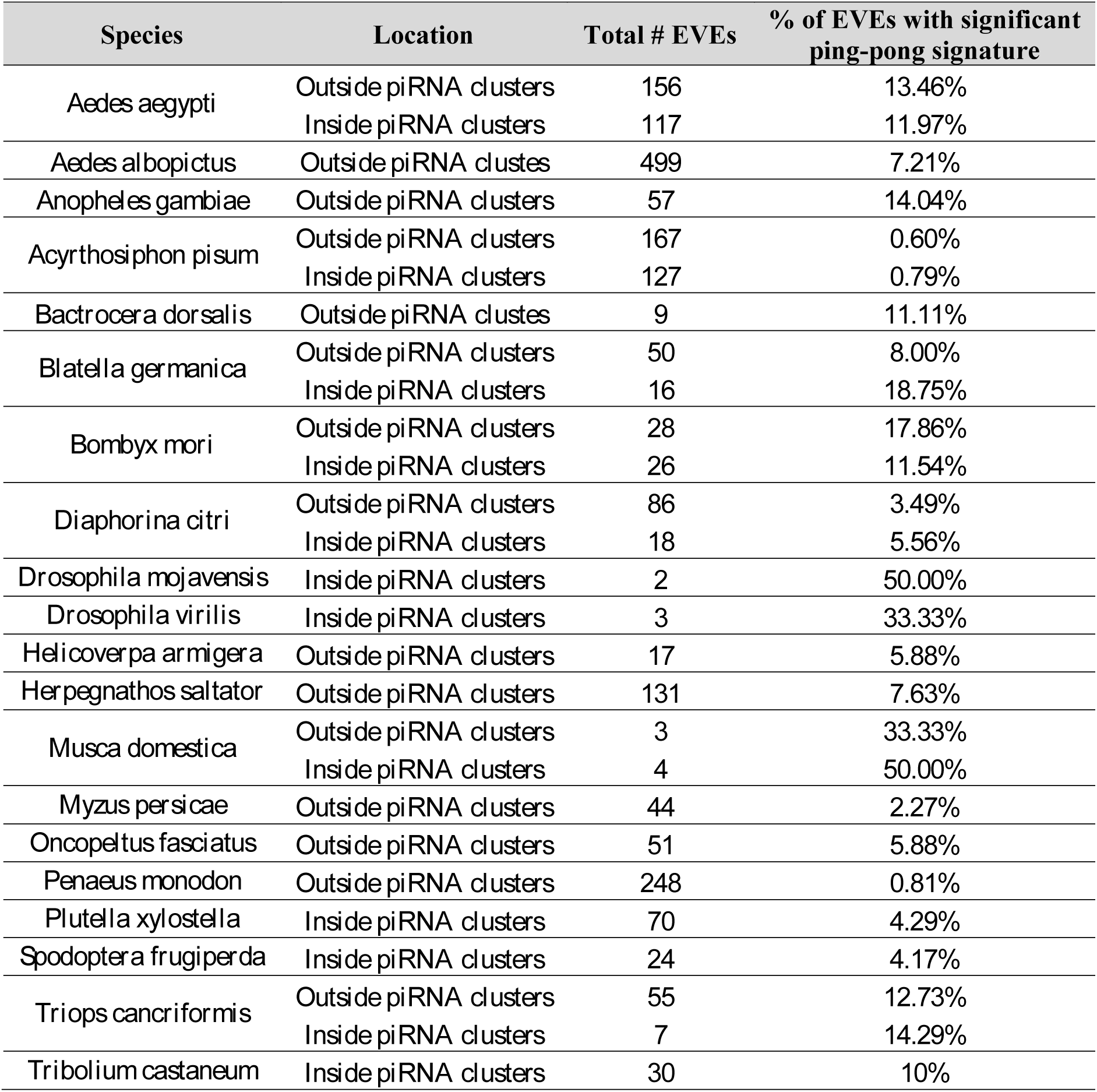
Percent of EVEs with mapped 24-32 nt sRNAs displaying a significant ping-pong signature

| Species | Location | Total # EVEs | % of EVEs with significant ping-pong signature |
| --- | --- | --- | --- |
| *Aedes aegypti* | Outside piRNA clusters | 156 | 13.46% |
|  | Inside piRNA clusters | 117 | 11.97% |
| *Aedes albopictus* | Outside piRNA clusters | 499 | 7.21% |
| *Anopheles gambiae* | Outside piRNA clusters | 57 | 14.04% |
| *Acyrtosiphon pisum* | Outside piRNA clusters | 167 | 0.60% |
|  | Inside piRNA clusters | 127 | 0.79% |
| *Bactrocera dorsalis* | Outside piRNA clusters | 9 | 11.11% |
| *Blatella germanica* | Outside piRNA clusters | 50 | 8.00% |
|  | Inside piRNA clusters | 16 | 18.75% |
| *Bombyx mori* | Outside piRNA clusters | 28 | 17.86% |
|  | Inside piRNA clusters | 26 | 11.54% |
| *Diaphorina citri* | Outside piRNA clusters | 86 | 3.49% |
|  | Inside piRNA clusters | 18 | 5.56% |
| *Drosophila mojavensis* | Inside piRNA clusters | 2 | 50.00% |
| *Drosophila virilis* | Inside piRNA clusters | 3 | 33.33% |
| *Helicoverpa armigera* | Outside piRNA clusters | 17 | 5.88% |
| *Herpegnathos saltator* | Outside piRNA clusters | 131 | 7.63% |
| *Musca domestica* | Outside piRNA clusters | 3 | 33.33% |
|  | Inside piRNA clusters | 4 | 50.00% |
| *Myzus persicae* | Outside piRNA clusters | 44 | 2.27% |
| *Oncopeltus fasciatus* | Outside piRNA clusters | 51 | 5.88% |
| *Penaeus monodon* | Outside piRNA clusters | 248 | 0.81% |
| *Plutella xylostella* | Inside piRNA clusters | 70 | 4.29% |
| *Spodoptera frugiperda* | Inside piRNA clusters | 24 | 4.17% |
| *Triops cancriformis* | Outside piRNA clusters | 55 | 12.73% |
|  | Inside piRNA clusters | 7 | 14.29% |
| *Tribolium castaneum* | Inside piRNA clusters | 30 | 10% |

## DISCUSSION

Mounting evidence points towards a role for EVEs in antiviral responses against corresponding viruses in animals and both transcription and translation of EVEs have been hypothesized to play important roles. Indeed, some EVEs possess features of purifying selection including maintenance of long open reading frames and low ratios of non-synonymous:synonymous mutations [30]. Moreover, experimental evidence indicating the functionality of EVE-encoded proteins has been shown in the thirteen-lined ground squirrel, the genome of which possess an EVE-encoded protein that inhibits replication of the corresponding virus *in vitro* [31]. Proposed mechanisms of transcription-mediated EVE-based immunity include the production of primary piRNAs from EVE-derived transcripts as well as the formation of dsRNA due to bi-directional transcription of EVEs and/or extensive secondary structure in EVE-derived transcripts [32].

Previous research indicates that EVEs are widespread in mosquito genomes and commonly produce piRNAs [14, 15, 18]. However, relatively little is known regarding the presence and functionality of EVEs in other arthropod species. Here we examined 48 arthropod genomes representing species belonging to 16 orders. We found that, as has been demonstrated in mosquitoes, EVEs are pervasive in the genomes of species spread throughout the arthropod lineage and frequently serve as templates for the biogenesis of piRNAs. Interestingly, we found that EVEs corresponding to negative sense ssRNA viruses comprised a plurality of the EVEs identified here. We also identified a large number of EVEs corresponding to viruses of the family *Parvoviridae*. As reported previously for *A. aegypti* and *A. albopictus*, we found that EVEs were enriched in piRNA clusters in a majority of species analyzed.

It has been proposed that EVE-derived piRNAs may play an antiviral role via the ping-pong cycle by directing post-transcriptional silencing of viral RNAs [15]. Cleavage of RNA targets by primary piRNA-guided argonaute proteins is dependent on base-pairing between primary-piRNAs and RNA targets [27]. However, unlike siRNA-directed cleavage, piRNA-directed cleavage appears to tolerate a small number of mismatches (∼ < 2-3) such that extensive, but not perfect, complementarity between piRNAs and their targets is required [27, 28]. While nucleotide identity between the majority of EVEs identified here and known viruses is generally too low to permit targeting of known viruses by EVE-derived piRNAs, 24-32 nt sRNAs mapping to 3.4% of EVEs possessed significant ping-pong signatures. These results raise the possibility that piRNAs derived from these EVEs may play roles in responses to infection with corresponding undescribed viruses.

We encountered a number of technical difficulties in our analysis. For some species, available genome assemblies and sRNA datasets were derived from different strains of the organism and in a small number of cases sRNA datasets derived only from one sex, only from particular organs, or only from certain life stages were available. These situations led to lower genome coverage of some species by mapped sRNAs, likely resulting in an underestimation of the number of EVEs producing primary piRNAs as well as the proportion of the genome annotated as piRNA clusters. Additionally, we found that when compared to experimental definitions of piRNA clusters, the piRNA clusters defined by proTRAC comprised smaller proportions of the genome. This may be due, in part, to the fact that the proTRAC algorithm was designed based on the characteristics of mammalian piRNA clusters, which display some important differences compared to arthropod piRNA clusters [10, 21]. Finally, the quality of genome assemblies in our analysis varied greatly. While the genome assemblies for some species such as *D. melanogaster* and *A. aegypti* are complete and well assembled, many genome assemblies are incomplete, highly fragmented, and contain duplications, particularly in repetitive regions such as piRNA clusters that typically contain a higher load of EVEs. Thus, we believe that as these genome assemblies improve, so too will our ability to accurately catalog the collection of EVEs present within them.

## CONCLUSIONS

An understanding of arthropod antiviral immunity is critical for the development of novel strategies to control vector-mediated virus transmission to animal and plant hosts. Our findings reveal that the important observations regarding the functionality of EVEs in mosquitoes apply to a wide range of other arthropod species and lend further support to the hypothesis that, in some circumstances, EVEs may constitute a form of heritable immunity against corresponding viruses. While EVEs may indeed occasionally provide the basis for an immunological response, we propose that given the lack of extensive nucleotide identity observed between EVEs identified here and currently described exogenous viruses, endogenization of viral sequences is an infrequent event and the ability of EVE-derived piRNAs to initiate a response against virus infection may decline over evolutionary time as exogenous viruses and their corresponding EVEs diverge. To gain an understanding of the general utility of the interaction between EVEs and the piRNA pathway as an antiviral mechanism, future studies should address the timescale over which acquisition of new EVEs takes places and to what extent genomic EVE content varies between geographically distinct populations of a given species.

## DECLARATIONS

### Authors’ contributions

AMH wrote the scripts and performed the analyses. JCN conceived the study, wrote the scripts, and performed the analyses. All authors analyzed the data, wrote the manuscript, and approved the final manuscript.

### Availability of data and material

The datasets and scripts used and/or analyzed during the current study are available from the corresponding author on reasonable request.

### Ethics approval and consent to participate

Not applicable

### Consent for publication

Not applicable

## Acknowledgements

No Applicable

## ADDITIONAL FILES

Additional file 1: GenBank accession numbers of genome assemblies. (XLSX 10 kb)

Additional file 2: GenBank Accession numbers of sRNA datasets. (XLSX 16 kb)

Additional file 3: Endogenous viral elements found within piRNA clusters in 48 arthropod genome assemblies via BLASTx. Species in which no Endogenous viral elements were found within piRNA clusters are not included. Species are separated into one species per sheet. (XLSX 103 kb)

Additional file 4: Endogenous viral elements found outside piRNA clusters in 48 arthropod genome assemblies via BLASTx. Species are separated into one species per sheet. (XLSX 409 kb)

Additional file 5: Viral families corresponding to endogenous viral elements found within piRNA clusters. (XLSX 13 kb)

Additional file 6: Viral families corresponding to endogenous viral elements found outside piRNA clusters. (XLSX 153 kb)

